# Blinks are strategically coupled with head movements in unconstrained natural gaze behavior

**DOI:** 10.64898/2026.06.29.731833

**Authors:** Alexander Goettker, Mary Hayhoe

## Abstract

Blinks are a ubiquitous yet largely unnoticed aspect of human vision, despite causing frequent interruptions of visual input that can amount to up to 10% of waking time. By leveraging a large dataset of unconstrained gaze behavior during two natural tasks, we found a novel behavioral strategy to limit the impact of blinks: blinks were strategically coupled with head movements, which minimizes information loss due to unreliable visual input during head movements. Specifically, blink probability increased with higher head velocities and showed strong temporal modulation relative to head movement onset: Blink probability was reduced before head movement initiation and then peaked during the head movement. The strength of this coupling was tailored to the individual needs of participants, with participants with higher baseline blink showing a stronger synchronization. This indicates that blinks are a part of an individually coordinated strategy when orchestrating eye and head movements during unconstrained natural behavior.

## Introduction

Although we blink every few seconds ^1,2^, we are rarely aware of it. This is notable, as every time we blink, we lose visual input, and given both the frequency and duration of blinking, it has been estimated that individuals can spend up to 10% of their waking time with their eyes closed ^3^. Researchers have long been fascinated by the question of why we do not notice these constant blackouts in our visual input ^4–6^. The critical factor seems to be an internal suppression mechanism that reduces visual sensitivity ^7,8^, which is similar to ideas for how we deal with the consequences of our own eye movements ^9,10^. However, the question remains: why are we not more affected by the constant loss of visual input in our everyday behavior?

Blinks are necessary to keep the cornea moisturized ^11,12^, which helps to improve the optical quality of the images ^13^. Thus, we cannot simply avoid blinking. However, there is growing evidence that blinks might also have other functions. Blinks might not be as detrimental to visual perception as previously thought, as visual performance after blinks can even be boosted (Ang & Maus, 2020), and this seems to be related to the visual transients on the retina caused by blinks ^15^. In addition, the timing of blinks also provides evidence for mechanisms that go beyond passive moisturization. We blink more frequently than would be necessary for moisturization ^13,16^, and when we blink is also not random: The timing of blinks is actively shaped by task demands ^17^: Hoppe and colleagues ^18^ showed that subjects rapidly learn how to time blinks to match the statistics of events they are required to detect. The timing of blinks can also be adapted to performing subsequent actions ^19–21^. In addition, when investigating isolated eye movements, previous studies found that blinks are more likely to occur during larger gaze shifts ^22–24^. Furthermore, recent work suggests that blinks can be related to cognitive processes, for example, to disengage from information ^25^, for example, when finishing a sentence or turning a page ^26,27^. Thus, within the constraints of specific tasks, blinks seem to be generally timed to minimize the risk of critical information loss.

However, how do we deal with the need to blink in less constrained, natural behavior ^28^? In typical vision experiments, participants perform a structured task with individual short trials, which determine when visual information is task-relevant. In addition, such experiments often also constrain movements by fixing the head, which is in stark contrast to the coordination of eye and head as a hallmark of natural gaze behavior ^29^. In the present study, we leverage a large-scale data set of unconstrained eye and head movements where participants perform two continuous natural tasks to reveal a strategy for blinking behavior that plausibly helps with the problem of minimizing information loss, namely, that blinks are coupled with head movements. This coupling shows a natural synchronization of blinks with longer times of less reliable visual input resulting from faster head movements. This strategy suggests that during unconstrained natural behavior, when to blink is part of an individually coordinated plan that orchestrates eye and head movements.

## Results

We used mobile eye tracking glasses (Pupil Labs Neon) to collect a large dataset of 81 participants completing two types of tasks: building blocks or copying a painting. Participants sat at a table and could move their eyes and heads without any constraints. The setup for both tasks was comparable: On one side of the table, there was either a manual that showed what needed to be built (see Figure 1A) or the original painting. On the other side of the table were either all available blocks or an empty sheet of paper and colored pencils. While completing these tasks, we used the eye tracker’s IMU to track head movements. By combining the head and eye-tracking data, we computed the gaze position in world coordinates (i.e., the eye + head position). Importantly, while participants completed these tasks, we recorded more than 38 thousand naturally occurring blinks and analyzed when they typically occurred.

**Figure 1.**
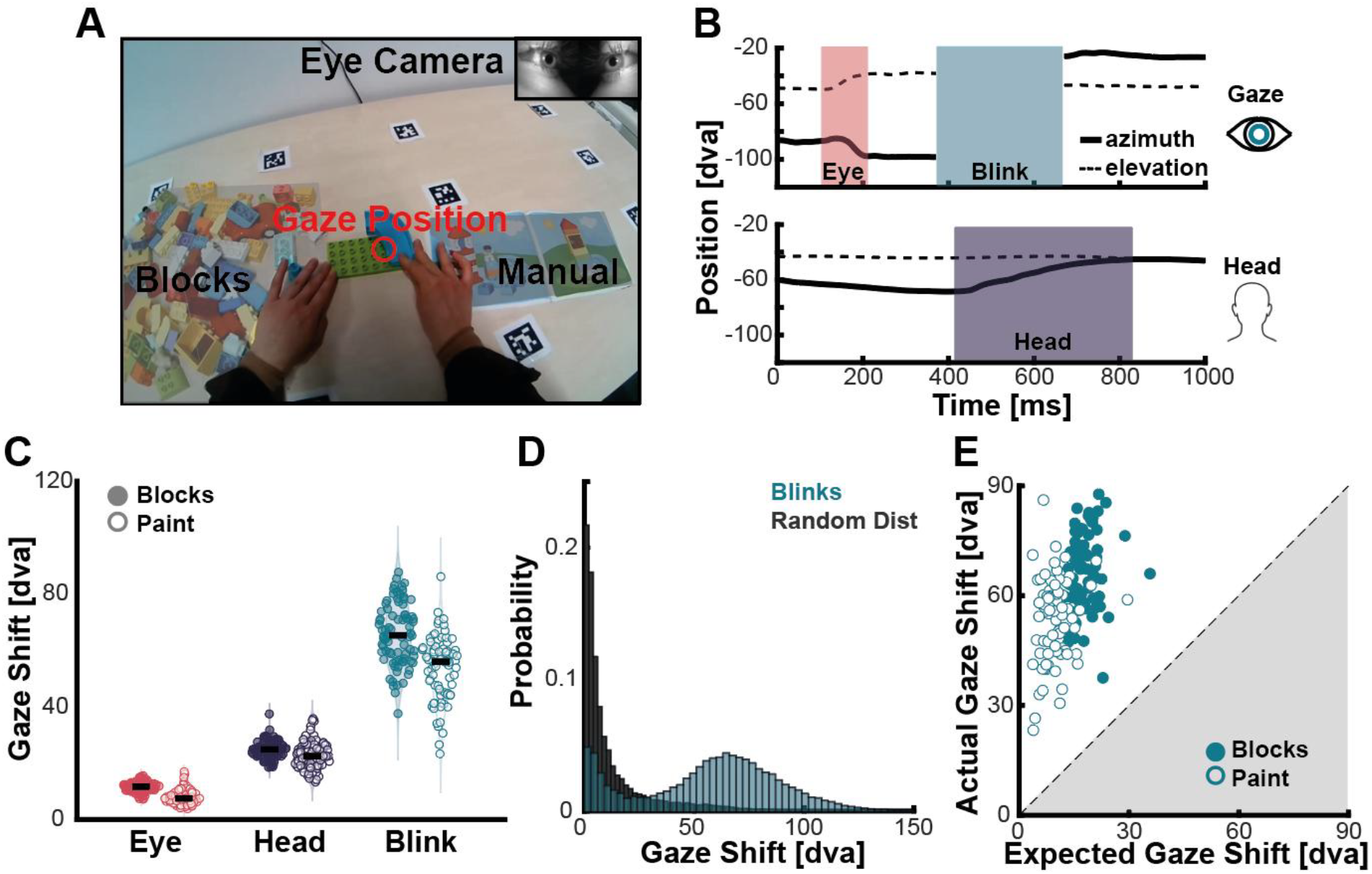
Gaze movements with unconstrained head. **A** An example image from the scene camera while performing the Blocks task. The red circle shows the gaze position, on the right side of the hands, the manual with the models that needed to be copied is shown. On the left side of the hands, all available blocks are visible on the table. The top right corner shows the image from the eye camera. **B** Raw behavioral data. The top panel shows the gaze position in world coordinates, the lower panel the head position. Solid lines show the azimuth, dashed lines the elevation. The highlighted colored areas are the phases where the respective movements have been detected. **C** The average gaze shift amplitude of eye movements (saccades), head movements, and across Blinks. Each dot depicts the average of one observer, the filled dots depict the Blocks task, and the open dots the Painting task. **D** The overall distribution of gaze shifts across >38k Blinks, in blue. The dark gray distribution is an estimate of the distribution if blinks were randomly distributed. **E** The expected gaze shift based on random blinking is plotted against the measured gaze shift across blinks. Each dot is one participant, the filled symbols represent the Blocks task, and the open symbols represent the Painting task.

### Blinks are coupled with large gaze shifts

When looking at the raw data (Figure 1B), we noticed a striking pattern: blinks often coincided with large gaze shifts. This also became evident when looking at summary statistics: the average gaze shift across blinks was much larger than the average gaze shifts caused by eye or head movements (see Figure 1C), sometimes even by several orders of magnitude. This indicates that blinks were frequently coupled with particularly large gaze shifts. To test for this coupling, we estimated an expected gaze shift distribution if blinks would occur at random times (Methods for more Details). When comparing the random with the actual distribution of gaze shifts (Figure 1D), you can already see that while the random distribution leads to mostly small gaze shifts, the distribution of the actual blinks has two peaks: one that follows the random distribution, but also a clear second peak, which is centered on large gaze shifts. Based on the random distribution, we could also directly compare the expected average gaze shift during blinks based on a random distribution with the observed mean gaze shift for each participant and each task (Figure 1E). The difference was extreme: Every subject in both tasks showed a higher actual mean gaze shift than what would have been expected from the random distribution (*t*(80)=41.425, *p*<.001, *d*=4.603 for Blocks, and *t*(80)=35.199, *p*<.001, *d*=3.911). It is important to note here that while gaze-evoked blinks for larger gaze shifts have been described for isolated movements ^22–24^, these results suggest that in unconstrained, sequential behavior, many blinks seem to be strategically coupled with very large gaze shifts.

### The occurrence of blinks is tuned to head velocity

After demonstrating this overall link between blinks and large gaze shifts, we tested whether we could establish a relationship between the characteristics of gaze shifts and the probability of blinking. Large gaze shifts in unconstrained behavior are typically combined shifts of eye and head (see Figure 1B), which means that the head is still moving during blinks. Since, by definition, we do not have access to eye movements during blinks, we focused our analysis on head movements. We hypothesized that high head velocities could be related to blinks. Fast head movements lead to large field visual motion on the retina, which makes the visual input less reliable and potentially challenges visual stability. To test this, we looked at all detected head movements (in total more than 109k). For each participant and task, we sorted the head movements by their peak velocity and then binned them in groups of 5% and computed the proportion of movements that contained a blink. Across observers and tasks, we found a lawful relationship between head velocity and the occurrence of a blink: the higher the head velocity, the higher the probability of blinking. Interestingly, this relationship was well described by a cumulative Gaussian, akin to a typical psychometric function (see Figure 3A for an example observer). This reveals that for large gaze shifts, there is a gradual and continuous scaling of the probability of blinking based on head velocity. This demonstrates a lawful relationship in the coupling of individual head movements to the likelihood of blinking, and highlights a strategic connection between faster head movements and blinks.

Although the overall relationship of more blinks during faster head movements was consistent across participants, the strength of the tuning of head velocity to blink rate differed on the individual level. As an indicator for the coupling of blinks and head movements for each individual, we looked at the P50-point of the fitted functions, the head velocity that led to blinks in half of the movements (see Figure 2B). While this velocity differed widely between participants, it was consistent between tasks (r=.798, p <.001), suggesting stable individual differences in the strength of the use of this strategy. Interestingly, we observed that especially participants with higher overall blink rates showed a stronger coupling between blinks and head velocities (reflected in lower velocities for 50% blinks, Blocks: r=-.795, p <.001; Painting: r=-.779, p <.001; see Figure 3C). This relationship seems to be particularly tuned to natural differences in blink rate, as these correlations stay highly significant even when controlling for the rate of head movements (Blocks: r=.-765, p <.001; Painting: r=-.727, p <.001). The same pattern was also true for the slope of the fitted functions: participants with higher blink rates showed a stronger increase in blink probability based on head velocity (Blocks: r=-.439, p<.001; Painting: r=-.594, p <.001). Together, this reveals that while not every blink is happening during a fast head movement (see also the first peak in Fig 1D), there is a strategic coupling of blinks to fast head movements. The strength of this coupling differs between participants, but especially participants who have an overall high blink rate, who almost certainly blink for head movements with a certain velocity (see Fig 2A).

**Figure 2.**
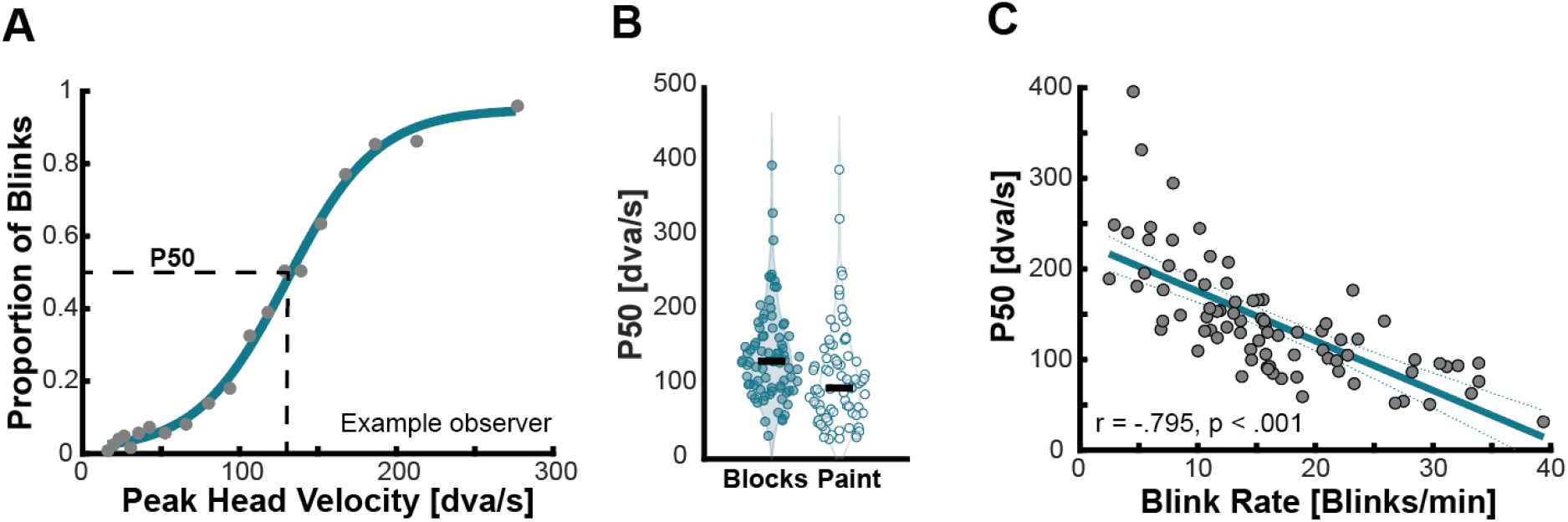
The tuning of blinks to head velocity. **A** The relationship between peak head velocity and blinking behavior for a representative participant. Data points show 5% bins of all head movements; the teal line shows a fitted cumulative Gaussian. **B** Distribution of the P50 of the fitted function for the Blocks and the Painting task. Each dot **C** The correlation between average blink rate per observer, and the 50% point of the fitted function in panel B. These data are for the Blocks task. Each dot shows one participant, the teal line a linear regression. Participants with higher blink rates show a stronger coupling of blinks and head movements.

**Figure 3.**
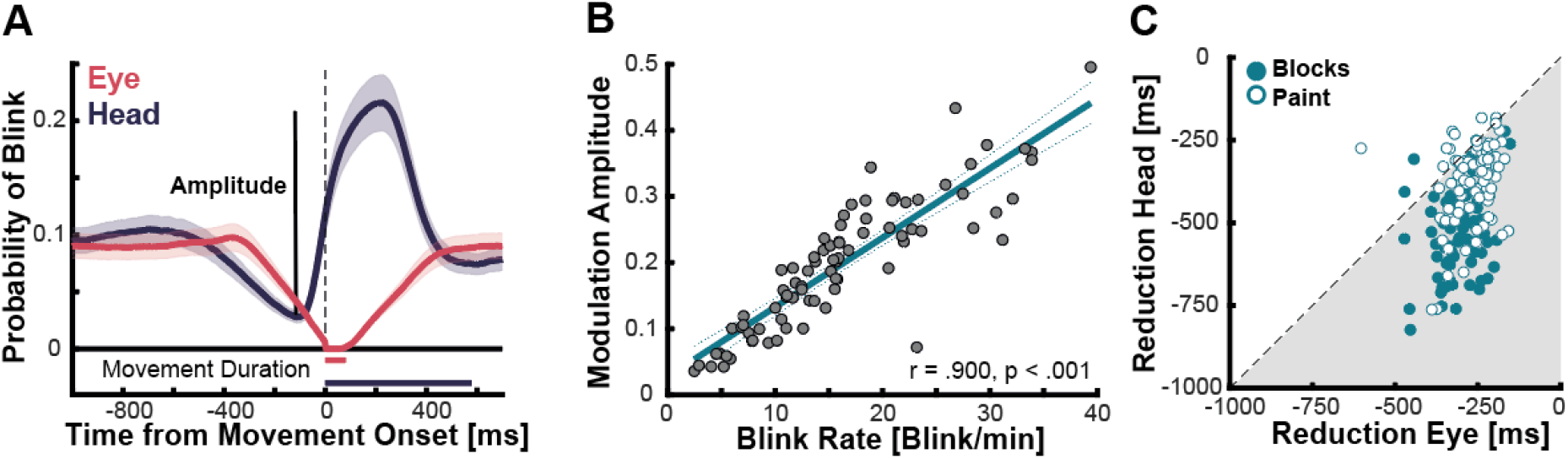
Temporal synchronization of blinks and head movements. **A** The time course of the probability to blink aligned to eye (blue curve) and head (red curve) movements. The line shows the average across observers for the Block task, the shaded area the 95% CI. The lines below zero depict the average movement duration. **B** The relation between the modulation amplitude of blink rate (see in Panel A) and the overall blink rate. Each data point depicts one observer for the Blocks task. **C** the comparison of the onset in the reduction of blink frequency for eye and head movements for the Blocks task (filled symbols) and the Painting task (open symbols). Each dot depicts one observer.

### Blinks are reduced before head movement onset and increased after movement onset

Our analysis so far revealed the tuning of the co-occurrence of blinks and head velocity. However, the big remaining question is whether the timing of blinks is actively adapted to match the timing of head movements. To test whether this is the case, we looked at the probability of a blink relative to head movement onset. We found a striking pattern (see Figure 3A, blue curve): Blink rate decreased beginning around 500 ms before head movement onset, reached a minimum shortly before head movement onset, and then peaked during the head movement and returned to the baseline rate afterwards. This suggests that blinks are actively suppressed before head movement onset, and instead, blink execution is indeed shifted to be temporally aligned with the head movement. In line with the results for the tuning to head velocity, we also observed that there is a strong relationship between individual differences in overall baseline blink rate and how strongly the blink frequency is modulated around the time of head movement onset (see Figure 3 B for Blocks, r=.878, p <.001 for Paint). This shows that participants with an overall higher need to blink not only show a stronger tuning to head velocity but also show a stronger temporal modulation of blink rate.

The peak in blink probability during the head movement demonstrates a systematic shift of blinks to co-occur with head movement. However, the initial reduction in blink rates before movement onset was not unique to head movements: when we performed the same analysis, but now centered it on eye movement onset (in total > 330k movements), we found a similar reduction in blink frequency (see Figure 3A, red curve). This reveals a more general reduction of blinks related to movement preparation. However, there was also an interesting difference in the initial reduction of blink probability between eye and head movements (compare red and blue curves in Figure 3A): We estimated the onset of the reduction per participant and task (see Methods for more Details), and it was significantly later for eye than for head movements (Blocks: t(79) = 14.944, p <.001, d = 1.671; Painting: t(79) = 8.365, p <.001, d = 0.935, Figure 3C). This difference suggests that there is a longer planning horizon for head movements than for eye movements, which may be a consequence of the higher motor costs and longer movement durations (see Figure 3A, bottom lines).

## Discussion

By leveraging a large data set of unconstrained eye and head movements, we revealed a naturally occurring strategic relationship between blinks and head movements. Blink frequency was tuned to head velocity and showed a strong temporal modulation with respect to head movement onset. Overall, the use of this strategy was also related to individual differences, with people with a higher blink rate showing a stronger coupling of blinks and head movements. These findings parsimoniously relate previous ideas regarding the active timing of blinks to minimize information loss and potential benefits of blinks for visual perception and show that during unconstrained natural behavior, blinks are integrated in the coordination of eye and head movements.

### Natural Blinking is temporally tuned to minimize information loss

While it is clear that we need to blink to moisturize the cornea ^11–13^, when we blink is strategically adapted to minimize information loss ^18,19^. Our recordings of natural blinking behavior revealed two novel behavioral strategies consistent with this idea.

First, blinks are strategically shifted to co-occur with head movements (see Figure 3A). This effect goes beyond previous research that demonstrated gaze-evoked blinking for single large gaze shifts ^22–24^, as it shows an active use of a strategy that ‘hides’ blinks at moments of unreliable visual input during head movements. The strategic shift of blinks into faster head movements is complementary to other results that show that there are special kinds of torsional eye movements that are only performed while blinking ^20^. To demonstrate the strategic relationship between head movements and blinks, we observed that blinking is not only temporally tuned to head movement onset, but also shaped by the upcoming head movement characteristics (Figure 2A). The coupling of blinking and head movements is tuned to head velocity, where especially fast movements lead to greater retinal motion. Fast head velocities (see Fig 2A) translate to similarly fast movements on the retina, as even when the eyes counterrotate to maintain fixation during fast head movements (the so-called vestibulo-ocular reflex, see ^30,31^), the peripheral visual input is still moving at velocities similar to head speed. Such high velocities, especially for the long durations of head movements, make the analysis of visual input difficult ^32,33^. The strategic link between the planning of head movements and blinks is also evident in results that show that for the same gaze shift amplitudes, blinks are more frequent when the head is involved ^23^ and that blinking activity is also registered when participants perform memory-guided head shifts while the eyes are closed ^24^. Thus, since blinking is generally necessary, performing blinks during fast head movements is clearly a good strategy, as it minimizes the loss of reliable visual information.

Second, we found a reduction of blinks before the execution of eye and head movements (Figure 3A). This shows that while accumulating visual information to plan the next movements, blinks are reduced to avoid interference with the ongoing visual target selection and to foster movement preparation. Such a reduction in blink rate mimics recent work that indicates that the reduction of small fixational eye movements (called “microsaccadic inhibition”) can serve as a proxy for the expectation of both sensory events ^34–36^, and self-planned movements ^37^. In addition, such temporal shifts of potentially interfering corrective saccades are not only present during fixation, but have also been reported when intercepting moving targets ^38^. Our results go beyond these previous findings by showing that such an inhibition of potentially interfering behavior (microsaccades or blinks), also generalizes to complex, sequential, and self-paced natural tasks. Thus, strategically adapting our behavior to minimize potential information loss and be most prepared for processing visual information at critical moments in time is a general mechanism that translates from artificial laboratory tasks to natural behavior.

### Natural blinking can benefit perception

The use of large head movements to shift gaze has a drastic effect. While eye movements can only change what can be fixated in an otherwise constant visual scene, head movements bring new parts of the world into the visual field. This makes the potential boost of visual processing after blinks especially beneficial ^14^. The critical aspect for this benefit seems to be the visual transients produced by the blink ^15^, and we observed that the blinks tend to be executed more towards the beginning of the head movement (see Figure 3A), allowing undisturbed processing of the visual information at the end of the gaze shift. This idea also fits well with the specific enhancement of low spatial frequencies ^15^. This helps particularly with coarse-to-fine analysis ^39^, which is especially critical when needing to process novel visual input.

In addition, the coupling of blinks and head movements could also serve a further purpose, namely, perceiving a stable world despite changes in retinal input resulting from self-motion ^40^. Work on visual sensitivity during eye movements has shown that a critical factor in not perceiving motion blur caused by our own eye movements is the masking of the blurred retinal image by the stable image present before and after the eye movement ^41^. However, while such a passive masking mechanism works for short eye movements, it is not feasible for longer head movements (typical saccade durations are 50-100 ms; the average head movement duration in our study is 550 ms). Motion blur in the middle of head movements would still be visible, so actively suppressing it by blinking might actually benefit the perception of a stable world. This fits well to the observation of higher blink frequencies when eye movements are performed over backgrounds that produce more blur ^22^. While it is important to note that participants also performed many head movements that did not include a blink, and subjects who generally do not blink much use head movements less to hide their blinks, across all participants and tasks, there was a lawful relationship between blink probability and head velocity (see Figure 2A). Instead of relying on the processing of a noisy, blurry image, especially during fast head movements, participants frequently used blinks to actively suppress it, and some participants even always blinked when the head moved above a certain velocity. Thus, while there are some similarities between the perceptual consequences of eye movements and blinks, there are also some clear dissimilarities (see ^6^). Thus, the percept during blinks is perhaps more similar to the mechanisms that create perception in the human blind spot. Here, we fill our visual experience based on some limited assumptions without actually having visual input ^42^. Similar to the lack of awareness of frequent blinks, subjects are quite confident in the accuracy of their perception in the blind spot, despite lacking actual visual input ^43,44^.

### Individuals strategically orchestrate blinks, eye and head movements to their advantage

Under natural circumstances, not every blink is related to a head movement (see the first peak of the gaze shift distribution in Figure 1C). Sometimes we have to blink to moisturize our eyes, even when no head movement is planned. However, our results demonstrate that people strategically plan and coordinate blinks, as well as eye and head movements together.

One particularly interesting aspect is the difference in the beginning of the reduction in blink rate before eye and head movements. This shows that blinks were inhibited in relation to the planning of the upcoming movement, and thus can be considered to reflect the planning horizon before a movement. The planning horizon was significantly longer for head movements (see Figure 3C), which makes sense as eye movements are considered to have relatively minimal costs due to their short duration and limited required effort ^45^. In contrast, head movements have longer latencies ^46^, take much more time (see Figure 3A), and due to the higher weight and inertia, are much more costly than eye movements ^45,47^. The differences in the planning horizon are also visible between the different tasks: the planning horizon for head movements is significantly longer in the block than in the paint task (Blocks: M=-531.27 ms, SD=138.21 ms; Paint: M=-384.57 ms, SD=124.95 ms, t(78) = 9.503, p <.001; see Figure 3C). This is directly related to the different roles that head movements have in these two tasks: in the painting task, head movements are mostly used to switch between the original painting and the piece of paper. In comparison, during the block task, participants frequently use head movements to scan the table to find fitting blocks. This more sequential, planned behavior is then reflected in a longer planning horizon for the block task.

The strategic coordination of blinks and head movements was also strengthened by the fact that it was used differently by our participants. There is converging evidence that individual differences can reflect an individual optimization of behavior: Participants prefer to look at different positions in faces, which is related to their own optimal point to discriminate them ^48^, can provide insights into idiosyncrasies in sensorimotor processing ^49–51^, or reflect a tuning of motor behavior towards one’s own cost functions ^52,53^. In line with these ideas, we observed that the more participants needed to blink, the stronger their coupling of blinks and head movements. This was true for the tuning to head velocity (Figure 2B), as well as the temporal modulation of blink rate (Figure 3B). In other words, the more often participants had to blink, the more likely it was that they embedded these blinks during head movements. This tuning is a good strategy as it helps them to address multiple problems with a single solution: they can moisturize their eyes by blinking, while at the same time minimizing the loss of critical visual information due to unreliable visual input, and potentially even getting benefits for visual perception.

Overall, our results highlight and emphasize the strategic control of blinks during unconstrained behavior in complex, sequential, and self-paced natural tasks. We show, in addition, that naturally occurring blinks and head movements are coupled, with blinks more likely during larger head movements. Individuals strategically orchestrate their behavior and include blink timing in movement planning. This pattern of behavior minimizes the loss of critical visual information, can benefit perceptual stability, and is tuned to the individual needs of a participant.

## Methods

### Participants

81 human observers (mean age 25.5 years; standard deviation: 4.46 years; 60 identified as female) completed the tasks. The recordings and the two natural tasks performed were part of a larger study, which was preregistered at https://osf.io/7utwn. All participants reported normal or corrected-to-normal vision (only possible with contact lenses due to the mobile glasses), and were naïve with respect to the study. Experimental procedures were in line with the Declaration of Helsinki and were approved by the Local Ethics Committee of the Department of Psychology and Sports Sciences of the Justus Liebig University Giessen (LEK-2024-0039). Written informed consent for recording the behavioral as well as the scene camera information was obtained from each participant. Participants received 8 Euros compensation per hour or course credits.

### Experimental setup and task

Participants sat on a chair in front of a table and could freely move. During the whole time, we recorded their eye and head movements as well as a scene camera video with the Pupil Neon Glasses (Pupil Labs, Berlin, Germany). Participants completed two tasks: *Blocks* and *Paint*. For the Blocks task, we used the Duplo Deluxe Brick Box (Number 10914), and the first four models in the manual had to be built. The setup was that the manual was on the right side of the table (inside a frame defined by some April Tags that were fixed on the table), and all available blocks were on the left side of the table. Then participants had to search for the fitting blocks, assemble the model, and once finished, disassemble it and start with the next model, until all models have been completed. In the Paint task, the setup was similar. However, instead of the manual, the model painting (two Mondrian patterns) was put on the right side of the table, and an empty piece of paper and colored pencils were on the left side of the table. The task was finished when the participants were happy with their paintings, and they completed one after the other.

### Preprocessing of behavioral data

The whole pipeline to label and extract the behavioral data is publicly available at: https://gitlab.com/AlexGoettker/labelmobiledata. To compute gaze shifts in real-world coordinates, we started by using the recording of the azimuth and elevation of the eye position (as eye in head position) and the IMU data to compute the gaze and head positions in world coordinates. For that, we followed the procedure available from [PupilLabs](https://docs.pupil-labs.com/alpha-lab/imu-transformations/). In short, we upsampled the IMU data from 110 Hz to 200 Hz, which is available for the eye position signal. We then used the quaternion values of the IMU to project them into a common reference frame. This procedure gives us the head azimuth and elevation, and the gaze azimuth and elevation (which is now eye + head). We then used a second-order Savitzky-Golay filter with a window size of 10 samples (50 ms) to filter these position traces. From the position traces, we compute gaze and head velocity by computing the distance between two consecutive samples on the sphere and multiplying it by the sampling frequency, which we then filtered with a second-order Butterworth filter with a cutoff frequency of 15 Hz.

### Blink Detection

We used the default blink detector by [PupilLabs](https://assets.pupil-labs.com/pdf/Pupil_Labs_Blink_Detector.pdf), and visually validated parts of the blink detection by checking whether, during fast head movements, the eye camera really shows a blink. Note here that while blinks are interpolated by default in the initial output data, we set the gaze position and velocity vectors during the blinks (+/- 5 samples or 25 ms around the blinks) to NaN.

### Labelling of Eye Shifts

Adapted from a similar approach by Niehorster et al. ^54^, we first use gaze velocity to identify slow and fast phases in the gaze movements as a preliminary step. To find slow phases, a sliding window approach was used. The size of the window was set to 1000 ms (or 200 samples). Within this window, the threshold for slow eye movements was set by an iterative procedure: the mean and the standard deviation of the velocity values were calculated, and then samples that were above the mean + 2.5 std of the velocity were excluded from this computation until there was no change in the mean estimate. The threshold for this window was then set to the final mean value + 2.5. std. Then the sliding window was moved by 10 ms (2 samples) and the procedure was repeated. This went through the whole recording,and for each point in time, the average threshold was computed across each iteration that this time point was within the sliding window. Then the slow phases were defined as periods during which the eye stayed below the respective adaptive thresholds consecutively for at least 80 ms. The epochs between those were considered as fast phases and our potential saccadic time points. To validate whether the fast phases actually could be considered saccades, we tested them based on the following criteria: (1) The peak velocity of the saccade had to be larger than 40 dva/s and smaller than 2000 dva/s; (2) the average velocity in the 25 ms before the labelled saccade and the average velocity in the 25 ms after the labelled saccade needed to be less than half of the peak velocity and the saccadic amplitude needed to be at least 0.1 dva. In addition, if the labelled fast phase was more than 100 ms long, we performed additional checks. To ensure that we have a consistent labelling of on and offset, we refined them by selecting the points, where the eye either only accelerated until the peak for the onset, and the last moment of deceleration after the peak for the offset. To check that we caught all saccades, we did a second pass, where, between the already successfully labelled saccades (+/- 40 ms), we identified periods where the gaze velocity was above 60 dva/s. Then, around the time the eye velocity crossed these thresholds, we again found potential saccade onsets and offsets and checked whether these saccades also fulfilled the above-mentioned criteria.

### Labelling of Head shifts

Head movements were defined based on head velocity. To qualify as a head movement, the head velocity needed to be above and stay above 10 dva/s or the 68^th^ Percentile of the head velocity for the next 10s for a minimum of 150 ms. If this was the case, then the first frame where the velocity was above the threshold was considered as the head movement onset, and the last frame above the threshold was considered as the head movement offset. After finding the onsets and offsets, we checked whether the so labelled head movement had a minimum amplitude of 1 dva, otherwise they were excluded. We also checked whether two consecutive head movements were at least 50 ms apart. If so both were considered valid; if not, and the difference in head movement direction was below 45°, then the two head movements were merged together. If they went into two different directions, we kept them as two separate head movements.

### Statistical Analysis

Gaze shifts were computed as the distance between the onset and offset in gaze position across all labelled eye and head shifts and across blinks. We then averaged the data per participant and task across all available movements. To estimate the expected gaze shift across blinks if they would occur randomly, we went through the data for each participant and task. We looked at the actual number of blinks and their duration and then randomly drew a starting point for each of them and matched their duration to the actual data. We then computed the gaze shift for these randomly drawn blinks and repeated this procedure 10 times per participant. To then estimate an expected blink gaze shift, we first averaged the data for each of the iterations across all blinks and then across iterations, and compared these values with paired t-tests to the actual mean gaze shift in the real data.

To test the relationship between head velocity and the likelihood of a blink, for each participant and task, we took all available head movements. We then sorted the trials into bins of 5% based on the peak head velocity. Then, for each group of such trials, we computed the probability that a blink occurred. We used this data to fit a cumulative Gaussian to the data (similar to a psychometric curve), and extracted two parameters: the head velocity that led to 50% of blinks (classically the PSE), and the standard deviation of the curve, which is related to the steepness of the curve (classically the JND). We excluded participants, where the estimated PSE was above 500 dva/s (this was the case for three participants in the Paint task). In a final step, we related the individual variations in these parameters across the two tasks, and also to some fundamental behavioral characteristics like the overall blink rate (defined as the number of blinks per minute), with Spearman correlations, or partial correlations controlling for differences in head movement rate.

To test for the temporal synchronization, we created a vector on a ms-level that contained a 1 when a blink occurred at this time, and a 0 when no blinks were performed. We then collected all eye or head movements for each participant and task, and saved the blink vector centered on movement onset from 1000 ms before the movement to 1000 ms after movement onset, we then took the average of these curves across all available movements, which lead to a vector per person and task that reflected the proportion of movements that contained a blink at each point in time. Afterwards, we used a moving average with a window size of 5 ms to smooth the time course. To visualize the data, we averaged these vectors across participants, but we also extracted two parameters from this time course per individual curve. First, we looked at the amplitude of the modulation of blink frequency for head movements as the difference between the maximum and the minimum of the curve. Second, to determine the onset of the initial reduction in blink frequency, we took each vector and subtracted the baseline frequency between 1000 and 500 ms before movement onset. We then searched the minimum of the vector before movement onset. From the minimum, we then went back in time to find the point when the vector for reached 90% of the reduction and when for the first time it reached 10% of the reduction. We then fitted a line to all the data between these two critical points and the onset of the reduction was then determined as the intercept of this line (as zero reflected the base rate). We excluded participants where this estimate was either 1500 ms before or after movement onset. This was the case for one participant in Blocks and one participant in Paint. We compared the extracted offsets with paired t-tests between eye and head movements.

## Author Contributions

Conceptualization & Methodology, A.G, M.H.; Software and Formal Analysis, A.G.; Writing – Original Draft, A.G.; Writing – Review & Editing, A.G., M.H.; Visualization, A.G.; Supervision and Funding Acquisition, A.G., M.H.

## Competing Interest Statement

The authors declare no conflict of interest.

## Data Availability

The raw gaze data will be publicly available at OSF. The analysis code will be made publicly available on GitHub.

## Acknowledgements

The authors want to thank Alexander Schütz, Ben de Haas and Karl Gegenfurtner for helpful discussions of the results and Svea Kürthen and Alicia Kaufmann for their help with the data collection. A.G. was supported by the Deutsche Forschungsgemeinschaft (Project No. 222641018–SFB/TRR 135 Project A1, Project No. 533717223–under Germany’s Excellence Strategy, EXC 3066/1 “The Adaptive Mind”). M.H were supported by NIH grant EY05729.

## Notes

### Competing Interest Statement

The authors have declared no competing interest.

